# Warm temperature impedes the spread of a heritable manipulative symbiont community in spider populations

**DOI:** 10.64898/2026.08.31.747884

**Authors:** Jennifer A. White, Jordyn D. Robinson, Matthew R. Doremus

**Affiliations:** S-225 Agriculture Science Center N, Department of Entomology, University of Kentucky, Lexington, Kentucky; 320 Morrill Hall, Department of Entomology, University of Illinois Urbana-Champaign, Urbana, Illinois

**Keywords:** co-infection, cytoplasmic incompatibility, feminization, heritable endosymbionts, *Rickettsiella*, thermal sensitivity, *Wolbachia*

## Abstract

Heritable bacterial symbionts are pervasive in terrestrial arthropods, often imposing reproductive manipulations to promote their own spread within host populations. Co-infections are common, potentially allowing symbiont co-infectors to hitchhike through a host population. However, adverse thermal conditions can disrupt these communities, particularly when co-infectors vary in their thermal sensitivity. We used a multi-generation experiment to test whether warm (29°C) conditions disrupted spread of heritable symbionts through uninfected populations of the spider, *Mermessus fradeorum*. We tested two common infection combinations: a single infection with a cytoplasmic incompatibility (CI) inducing *Rickettsiella* or a feminizing co-infection that included a feminizing *Wolbachia*, the same *Rickettsiella*, and up to three apparent hitchhikers (two additional *Wolbachia* strains and *Tisiphia*). We initiated replicate populations with 1/3 of one infection type and 2/3 uninfected spiders, evaluating population infection rate over 5 spider generations under different temperature regimes. Under cool (21°C) conditions, *Wolbachia* feminization drove co-infection to 88% and *Rickettsiella* CI drove single infection to 83% of host populations. Vertical transmission for all symbionts was high (97-99%) and hitchhiking symbionts also spread effectively. Under warm conditions, feminization and CI efficacy were reduced, and symbionts suffered variably reduced vertical transmission. Warm conditions ultimately destroyed the co-infecting symbiont consortium and impeded symbiont spread. On its own, though, *Rickettsiella* was still able to increase, despite reduced strength of CI. We hypothesize that contrasting tensions between feminizing spread of the symbiont consortium versus environmentally driven loss of function and transmission may explain observed patterns of mixed infections in field populations of this spider.

## Introduction

Heritable bacterial symbionts are pervasive in terrestrial arthropods (Weinert et al. 2015). Passed from mother to offspring, often within the cytoplasm of the egg, these microbes are typically highly dependent on their host, having often lost the ability to live independently and lacking frequent opportunity to shift horizontally among host lineages (Brown et al. 2020; Douglas 1998; Moran et al. 2008). While such dependence selects for minimizing negative effects on their host because of their shared fate, maternal inheritance conversely sets up a potential reproductive conflict between symbiont and host. Many maternally inherited microbes have evolved mechanisms to manipulate their hosts’ reproduction to favor their own transmission by increasing the production and/or fitness of female offspring (Engelstädter and Hurst 2009; O’Neill et al. 1997). These manipulations include induction of parthenogenesis, male killing, feminization of genetic males into functional females, distortion of female sex allocation choices, and cytoplasmic incompatibility (CI), which sabotages matings between uninfected females and infected males (Brenninger et al. 2025; Werren et al. 2008). Such manipulations allow maternally inherited symbionts to spread and persist in host populations, even if they impose a cost upon their host (Turelli 1994). Heritable symbionts that manipulate reproduction are globally widespread, but geographic patterns (Ahmed et al. 2016; Charlesworth et al. 2019; Hague et al. 2022) suggest underlying climatic factors may drive the distribution of symbionts, with potentially important consequences for the biology, evolution, and ecology of their arthropod hosts (Corbin et al. 2017; Engelstädter and Hurst 2009).

Temperature is increasingly recognized as an important abiotic driver of symbiont biology. Many heritable symbionts have thermal optima (Anbutsu et al. 2008; Corbin et al. 2017; Doremus et al. 2019), decreasing in efficacy as temperatures increase (Chrostek et al. 2021; Nasehi et al. 2022), or decrease (Corbin et al. 2021; Hague et al. 2022; Hague et al. 2024). Warm temperatures, in particular, have frequently been shown to decrease the penetrance of symbiont-induced phenotypes and/or transmission efficiency (reviewed in Martins et al. 2023), but often in a strain-or host-specific manner (e.g., Bagchi et al. 2026; Ross et al. 2017). Field studies have indicated that use of cytoplasmic incompatibility (CI) to drive symbionts through target vector populations can fail under thermal extremes (Ross et al. 2019), demonstrating the importance of understanding the nuances of symbiont-host interactions under different temperature regimes. Such interactions might be particularly complex when multiple symbionts, with different thermal profiles, inhabit the same host.

Co-infection by multiple strains of heritable symbionts is quite common in arthropods, with certain host clades (e.g., whiteflies, aphids, spiders) being particularly prone to multiple infections within the same host individual (Rock et al. 2018; Skaljac et al. 2010; White et al. 2020). Disentangling the various roles these microbial community members play within their shared host can be complex, but it appears they variously contribute facultative benefits to their hosts (Corbin et al. 2017; Oliver and Higashi 2019), induce contrasting reproductive manipulations (Montenegro et al. 2006; Yoshida et al. 2019), or sometimes are simply parasites or commensals, hitchhiking from host generation to generation of host thanks to the effects caused by their co-infecting colleagues (Doremus and Oliver 2017; Rock et al. 2018). Longstanding consortia may have the opportunity to evolve complementarity of function and minimize costs to the host (Engelstädter et al. 2007; Vautrin and Vavre 2009), but environmental disruption has the potential to introduce dysfunction within the community (Mackevicius-Dubickaja et al. 2026).

The spider *Mermessus fradeorum* (Linyphiidae) can host up to five co-infecting strains of endosymbiotic bacteria that impose different modes of reproductive manipulation: a *Rickettsiella* (R) that causes CI (Rosenwald et al. 2020), a strain of *Wolbachia* (W1 or *Wolbachia* 1) that feminizes genetic males into functional females (Curry et al. 2015; Mackevicius-Dubickaja et al. 2025), and a *Tisiphia* (T) and two additional strains of *Wolbachia* (W2, W3) that have unknown effects on the host, although they appear to strengthen feminization caused by *Wolbachia* 1 (Mackevicius-Dubickaja et al. 2025). Whether these other symbionts cause direct phenotypic effects on the host or not, they may still persist and spread in the host population as co-infecting hitchhikers with their manipulative colleagues (Doremus and Oliver 2017; Hurst and Jiggins 2005; Rasgon et al. 2006; Schuler et al. 2016). Local field populations of the spider are almost universally infected with one or more of these symbionts: almost all spiders are infected with *Rickettsiella*, with approximately half exhibiting co-infections by one or more of the other symbionts (Rosenwald 2020; Rosenwald et al. 2020; J. White unpublished data). Co-infections that include the feminizing *Wolbachia* 1 represent approximately 20% of sampled local populations, but are less prevalent in some other geographic areas (Rosenwald 2020; Rosenwald et al. 2020; J. White unpublished data). Most spiders that have *Wolbachia* 1 have the full consortium of all 5 symbionts.

Temperature affects all of the symbionts within *M. fradeorum*, but in different ways and to different degrees. Both the strength of CI and feminization weaken with increased temperature (Mackevicius-Dubickaja et al. 2026; Proctor et al. 2024), but at different timepoints. Spider development at warm temperatures (26-29°C) directly reduces *Rickettsiella*’s ability to induce CI in warm-reared males (Proctor et al. 2024), but has a time delayed effect on feminization, reducing *Wolbachia* 1’s ability to feminize the offspring of heat-treated females (Mackevicius-Dubickaja et al. 2026). Further, warm temperatures also reduces vertical transmission of all the symbionts except *Rickettsiella*, with *Tisiphia* being particularly sensitive (Mackevicius-Dubickaja et al. 2026). Thus, warm temperature may disrupt co-infecting consortia, with the potential to hinder the ability of the symbionts to persist and spread within host populations.

Here, we directly test the effect of temperature on symbiont spread within laboratory populations of *M. fradeorum*. We initiated replicate populations of spiders that were composed of a mixture of infected and uninfected individuals, and asked whether the proportion infected changed over time as a function of temperature. We did this both with combinations (“symbiotypes”) of co-infecting symbionts that included the feminizing *Wolbachia 1*, and separately with spiders infected only with the CI symbiont, *Rickettsiella*. Our study fills a critical gap in the literature, bridging between short-term laboratory experiments that document clear effects of temperature on symbiont-induced phenotypes (Doremus et al. 2018; Hague et al. 2022; Higashi et al. 2020; Proctor et al. 2024; Ross et al. 2017) but lack longitudinal population tracking, and field–based syntheses that show geographic or temporal patterns (e.g. Charlesworth et al. 2019; Zchori-Fein et al. 2014), but have covarying complexities that preclude the ability to determine causality. We find that under cool conducive conditions, both the feminizing consortia and the CI symbiont spread in uninfected laboratory populations, with some populations reaching symbiont fixation within 5 generations. Under warm conditions, the feminizing consortium was disrupted, spawning a variety of subsidiary symbiotypes, with the frequency of the feminizing symbiont and most other symbionts decreasing over generations. The CI symbiont was able to effectively spread through populations at warm temperatures, but only when it was the sole symbiont in the population. In co-infection contexts, the CI symbiont failed to appreciably increase in prevalence in warm conditions, instead maintaining a stable infection rate across five spider generations. We therefore hypothesize that observed field population mixtures of spiders with the feminizing consortium versus only the CI symbiont may result from contrasting tensions between feminizing spread of the consortia and environmentally driven loss of function and transmission of most of the symbionts.

## Methods

### Does temperature affect spread of feminization

We tested the ability of the feminizing consortium to spread in uninfected host populations under different temperature regimes using 4 replicate populations per temperature. We initiated each population with 30 female and 30 male spiders, individually housed in condiment cups (4cm diameter) with moistened plaster at the bottom for humidity control. Ten of the initial female spiders per population contained feminizing consortia as one of three specific symbiotypes: all 10 were infected with *Rickettsiella*, *Tisiphia*, and the feminizing *Wolbachia* 1 (RTW1), but subsets additionally contained *Wolbachia* 2 (RTW12), or both *Wolbachia* 2 and 3 (RTW123). Each replicate population started with approximately the same proportion of the three feminizing symbiotypes. The remaining 20 female spiders and all male spiders were uninfected. All initial spiders had been reared under the same conditions (20°C constant temperature, dark conditions, fed twice weekly) and originated from the same spider matrilines (3 feminized, 5 uninfected), although spider age at mating was widely variable in this F0 generation. Uninfected lines had been previously cured of facultative symbionts with antibiotic treatment (Rosenwald et al. 2020) and had been maintained as uninfected lineages for multiple generations prior to experimental use.

For each population, we randomized mating pairs of male and female spiders and fed each spider one fruit fly prior to being paired for 4 hours to mate. Following mating, female spiders destined for warm populations were moved to an environmental chamber maintained at 29°C whereas cool population females were moved to an environmental chamber maintained at 21°C. All females were fed 1-2 times and allowed up to one week to lay one eggmass, then they were deprived of food for a minimum of 3 days before being preserved in 95% ethanol. Males were deprived of food and preserved directly after mating. Using this protocol, the vast majority of spiders effectively mated and laid egg masses, but a subset of spider pairings per population either failed to mate or failed to propagate (mean = 8.2% per population per generation), reducing the effective population size per generation. Such failures proved to be random with respect to maternal infection, cross type, and temperature, and therefore simply were viewed as part of the random events experienced by each population.

Each egg mass cup was provisioned with collembola (*Sinella curviseta*) prior to spiderling hatch to diminish cannibalism among siblings. Once hatched, we counted spiderlings and dissected the egg mass to count any remaining unhatched eggs. We transferred each hatched spiderling to an individual cup with its own supply of collembola; cool populations averaged (mean ±S.E.) 384 ±13 separated spiderlings per population per generation, and warm populations averaged 350 ±14, due to smaller size and faster metabolism experienced at warmer temperatures. Older spiderlings were fed one wingless *Drosophila melanogaster* fly twice weekly. Once 95% of spiderlings were large enough to determine sex based on palp development (sex can be determined two instars before adulthood), female and males were separately randomized, and 30 females and 30 males were selected to initiate the next generation of the population. An additional randomized 15 females and 15 males were reserved as extras, to be substituted in case of mishap, mortality, or misallocated sex of the selected 30. The remaining cohort of spiderlings was discarded. Selected spiders were maintained with twice weekly feedings until the next generation was propagated. Warm populations were propagated on an 8-week cycle, cool populations on a 12-week cycle. All populations from the same temperature regime were offset from one another on two-week intervals for logistical reasons, and each population was maintained through the mating of generation F5.

Because we reared spiders individually, we tracked the pedigree and expected infection status of each spider in the experiment. For spiders that were expected to be infected, we confirmed the presence of all symbionts via diagnostic PCR (see below). For spiders that were expected to be uninfected, we checked all F0 and F5 females, and also checked if a supposedly uninfected spider lineage showed potential evidence of infection (female-biased sex ratios or potential CI).

We analyzed the population spread of each of the five symbionts associated with the feminization consortium (*Rickettsiella*, *Tisiphia*, *Wolbachia* 1-3) using a binomial generalized linear mixed model (GLMM) with a logit link function. In each model, the response variable was symbiont presence, and the model included generation (F0-F5) and temperature (21°C, 29°C) as fixed effects with an interaction. Population was included as a random effect to account for overdispersion. We calculated estimated marginal means for each generation × temperature contrast to compare predicted probabilities of infection with each symbiont.

We next analyzed major factors potentially driving feminizing symbiont spread: sex ratio, offspring production, and hatch rate. We analyzed offspring production and hatch for generations F1-F5, but did not raise the final generation of offspring to adulthood, and thus only analyzed offspring sex ratio for generations F1-F4. The initial F0 generation was excluded from sex ratio and offspring production analyses because F0 spiders were not reared under the different temperature treatments. We analyzed these data two ways. First, we categorized mothers as either uninfected with any endosymbiont (uninfected) or infected with any combination of symbionts (infected) and compared the sex ratio of their offspring at both temperatures using a binomial GLMM with female offspring per brood as the response variable. Temperature (21°C, 29°C), maternal infection status (infected, uninfected), and their interaction were fixed variables, while population was included as a random effect. We similarly analyzed offspring production as the total number of spawn produced by each female using a negative binomial GLMM with the same factors. We calculated estimated marginal means for each temperature × infection contrast to compare predicted probabilities of female offspring production and egg counts. Second, due to inconsistent vertical transmission and symbiont loss in the warm populations, we performed additional sex ratio and egg production analyses using an infected category modified to strictly include those spiders that retained the feminizing *Wolbachia* 1 symbiont. Uninfected spiders were expanded to include spiders with unfeminized symbiotypes lacking *Wolbachia* 1 (e.g. R-only, RW2, RW23, etc). All other model parameters were identical to the original sex ratio and spawn production models.

Because feminized populations occasionally produced infected males, we also evaluated the strength of cytoplasmic incompatibility across temperature treatments. We analyzed the number of unhatched eggs as a proxy for offspring survival across the four possible cross types (female ×male): uninfected × uninfected, uninfected × infected (incompatible CI cross), infected × uninfected, and infected × infected. We used a zero-inflated negative binomial GLMM with unhatched eggs as a response variable and temperature (21°C, 29°C), cross type (four types), and their interaction as fixed effects. Population was included as a random effect. We calculated estimated marginal means for each temperature × cross type contrast to compare predicted probabilities of offspring mortality. We used glmmTMB v.1.1.14 (Brooks et al. 2017) to fit all GLMM models and assessed significance of fixed effects on symbiont spread, offspring sex ratio, and offspring mortality using Type II ANOVA with car v.3.1.5 (Fox and Weisberg 2018). We tested model fit using DHARMa v.0.4.7 (Hartig 2024) using tests for dispersion, outliers, Kolmogorov-Smirnov test for residual normality, and zero-inflation. We calculated estimated marginal means for all contrasts using emmeans v.2.0.3 with type = “response” (Lenth and Piaskowski 2026). P-values and 0.95 confidence intervals used the Tukey correction method for multiple comparisons. All GLMM statistical tests were run in R v.4.5.1 (R Core Team, 2025).

### Vertical transmission

We used diagnostic PCR to track the infection status of the majority (1074/1109 = 96.8%) of the propagated infected spiders within the feminization populations. Each specimen was extracted using DNeasy (Qiagen) extraction kits according to manufacturer’s protocol, then screened using symbiont specific PCR primers and reaction conditions for all 5 symbionts (**Supplemental Table 1**) as described in Mackevicius-Dubickaja et al. (2025). For specimens that were negative for all symbionts, we also ran diagnostic PCRs for the CO1 gene (Folmer et al. 1994) to check extraction integrity. Any specimens that failed to amplify CO1 product were excluded from further analysis.

We then compared symbiotype of mother and offspring from generations F1-F5. In all instances where offspring appeared to gain a symbiont that was not detected in their mother, we re-ran the PCR for both mother and offspring. If the discrepancy persisted (4 cases/864 = 0.4%, representing 3 mothers), we assumed the cause was either a poor extraction (or low titer infection) in the mother, or a contamination event where the offspring was misassigned to the wrong mother. We excluded these cases from vertical transmission calculations, but used them to set expectations for error rate in instances of symbiont loss between mother and offspring. Because instances of symbiont loss in offspring were much more common (especially in warm populations) we did not re-run all PCRs that indicated loss of a symbiont between mother and offspring, but we did double check all apparent cool population losses and a subset of apparent warm population losses, especially where siblings differed in their symbiotype. We used logistic regression with Williams’ correction for moderate overdispersion (Arc v1.06; Williams 1982) to compare vertical transmission rates among symbionts within each temperature regime, and for each symbiont in cool versus warm conditions, excluding the F0 generation as in the sex ratio and offspring production analyses.

### Does temperature affect spread of CI

We conducted a CI spread experiment simultaneously with the feminization spread experiment, using parallel methodology except as follows. The infected portions of the F0 generation were initiated with matrilines that were infected with only the CI-inducing symbiont *Rickettsiella* (7 matrilines). One third of both females and males were *Rickettsiella*-infected in the F0 generation. We again started with 4 cool and 4 warm populations, but one of the cool populations was accidentally frozen just after the mating of the F3 generation, when no replacement spiders from the same population were available. We reconstructed a population of exactly the same infection status and cross-types of the lost population using leftover spiders from one of the other populations, and carried on with this substitute population for the remaining generations. We also initiated an additional cool population that trailed the rest of the experiment, but was initiated from F0 exactly like the rest of the populations. Consequently, we had 5 cool CI populations and 4 warm CI populations. Because *Rickettsiella* transmission is much more reliable than the other symbionts even under warm conditions (Mackevicius-Dubickaja et al. 2026), we did not double check the infection of every infected individual, but instead checked for *Rickettsiella* infection of females in the first and last generation. In several instances in warm populations where *Rickettsiella* had apparently been lost by F5, we backtracked through previous generations to determine where the loss (or misallocation of a spiderling) occurred, and reclassified descendant infection status accordingly.

As with the feminizing cohort, we analyzed the spread of *Rickettsiella*, as well as F1-F4 offspring sex ratio, and F1-F5 offspring production and CI mortality using a series of GLMMs in R. We analyzed the spread of CI *Rickettsiella* through uninfected populations at two temperatures (21°C, 29°C) using a binomial GLMM with *Rickettsiella* presence as the response variable and generation (F0-F5), temperature, and their interaction as fixed variables. We additionally analyzed offspring sex ratio using a binomial GLMM with female offspring per brood as the response and temperature, maternal infection, and their interaction as fixed variables. To evaluate strength of CI, we compared offspring mortality in egg masses using a zero-inflated negative binomial GLMM with unhatched eggs as the response and temperature, cross type, and their interaction as fixed variables. We next evaluated fecundity using a negative binomial GLMM with total spawn produced as the response and temperature, female infection, and their interaction as fixed variables. We included population as a random effect to account for overdispersion for all GLMMs and tested significance of fixed effects using ANOVA. We calculated estimated marginal means for contrasts, using the Tukey method of p-value correction to account for multiple comparisons.

## Results

### Warm temperatures inhibit spread of feminizing symbiotypes within a population

Qualitatively, we found that feminizing symbiont infection did not spread as efficiently in warm populations as cool populations. Under cool conditions, 88.3±7.2% of the female population was infected by F5, with feminizing symbiotypes reaching fixation in two out of four populations. In all cool populations, the percentage of infected females more than doubled from the starting percentage of 33.3%. In contrast, in warm populations only 49.1±12.5% of females were infected by F5, and infection did not reach fixation in any warm populations. Infected females constituted 76.7% of the best performing warm population, but had decreased to only 16.7% in the worst performing population (**Figure 1**).

**Figure 1.**
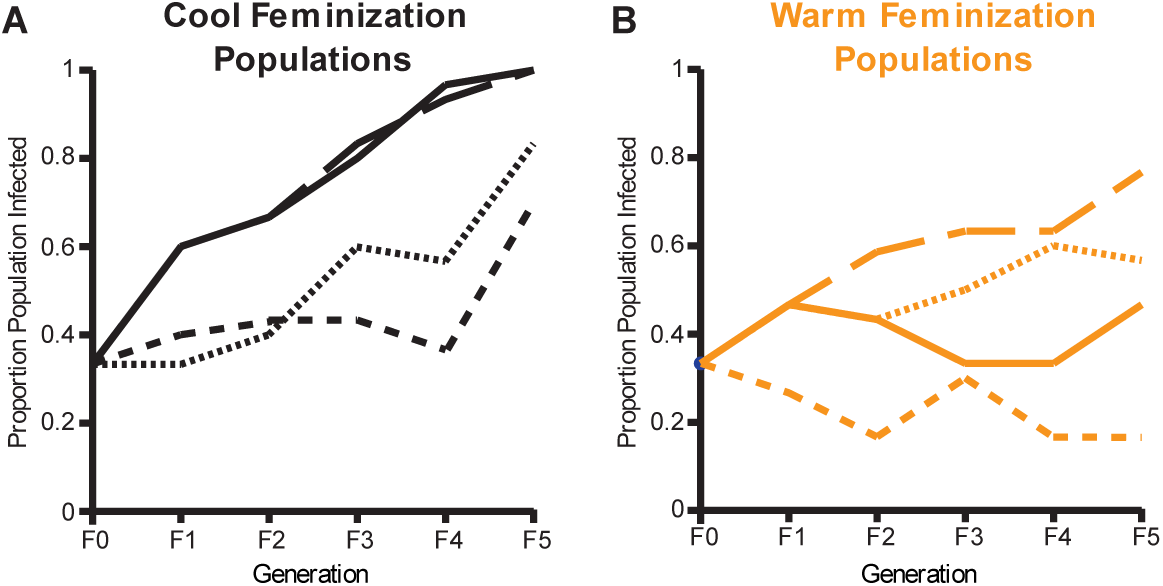
Proportion infection of laboratory *Mermessus fradeorum* populations under A) cool (21°C) or B) warm (29°C) conditions in the feminization spread populations. Each line represents a replicate population, with proportion infection depicted for the 30 adult females that were randomly selected to propagate the next generation. Infected spiders were infected with any combination of *Rickettsiella, Tisiphia,* and *wolbachia* strains 1, 2, and 3.

Further, the “infected” portion of these warm populations rarely included the feminizing symbiont, *Wolbachia* 1, or reflected the starting symbiotype due to reduced vertical transmission efficiency. Under cool conditions, all five symbionts had very high transmission rates throughout the experiment (97.0-99.7% transmission per generation across the symbionts; **Figure 2**), and did not differ significantly among symbionts (ΔDev = 1.17, d.f. = 4, P = 0.883). Under warm conditions, the CI-causing symbiont, *Rickettsiella*, still enjoyed very high transmission rates (97.1±0.6% per generation), but was significantly lower than the near perfect transmission (99.7±0.3%) under cool conditions (ΔDev = 11.8, d.f. = 1, P < 0.001). The other symbionts experienced more substantial reductions in vertical transmission in warm versus cool populations. Consistent with previous findings (Mackevicius-Dubickaja et al. 2026), *Tisiphia* transmission failed entirely from F1 to F2 in warm populations, leading to extinction of this symbiont from all four warm populations. For the three *Wolbachia* strains, vertical transmission ranged from 40-80% per population per generation. All three *Wolbachia* strains experienced significantly lower transmission in warm than cool conditions (W1 ΔDev = 29.2, d.f. = 1, P < 0.001; W2 ΔDev = 76.3, d.f. = 1, P <0.001; W3 ΔDev = 13.1, d.f. = 1, P < 0.001). The net effect of these progressive transmission failures was that by the end of the experiment, most of the infected spiders in warm populations only retained the CI symbiont, *Rickettsiella* (**Figure 3**, **Supplemental Figure S1**). *Wolbachia 2* was nearly extinct along with *Tisiphia*, and the feminizing symbiont *Wolbachia 1* was present in only 22.1 ± 9.1% of the infected spiders (**Figure 3**, **Supplemental Figure S1**). In contrast, in the cool populations, the distribution of symbiotypes among infected spiders remained virtually identical throughout the experiment, and all infected spiders in F5 retained the feminizing *Wolbachia1* (**Figure 3**, **Supplemental Figure S1**).

**Figure 2.**
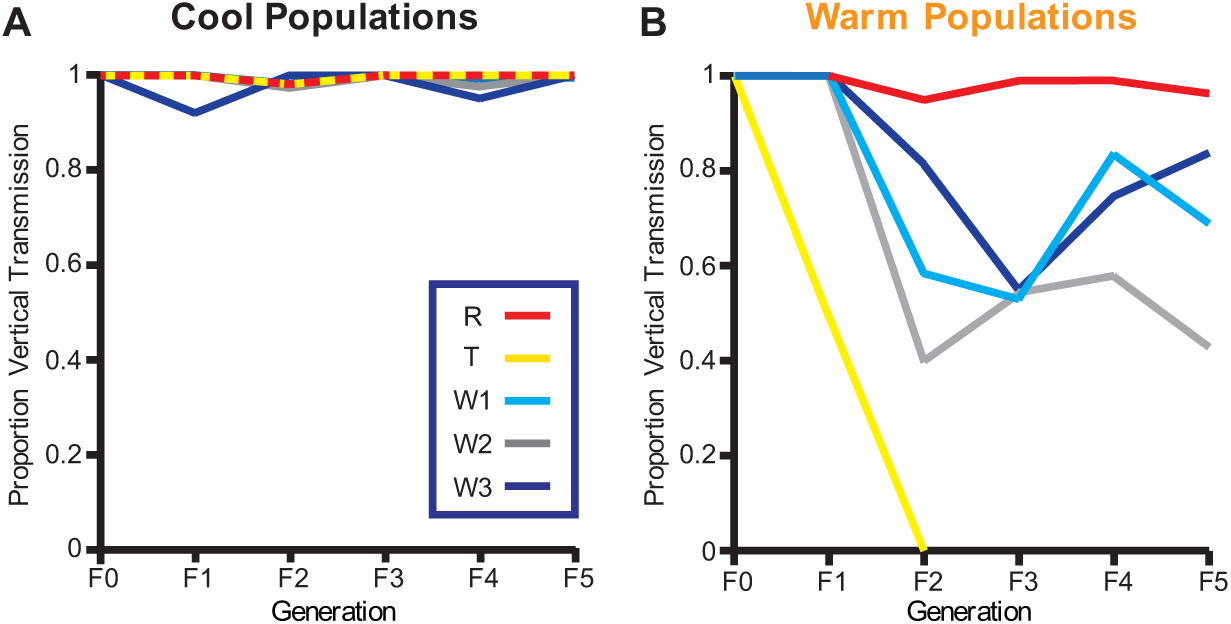
Vertical transmission of five bacterial symbionts in laboratory *Mermessus fradeorum* populations under A) cool (21°C) or B) warm (29 °C) conditions. Each line represents a different symbiont: R= *Rickettsiella,* T *= Tisiphia,* W1 = *Wolbachia* 1, W2 = *Wolbachia* 2, W3= *Wolbachia* 3.

**Figure 3.**
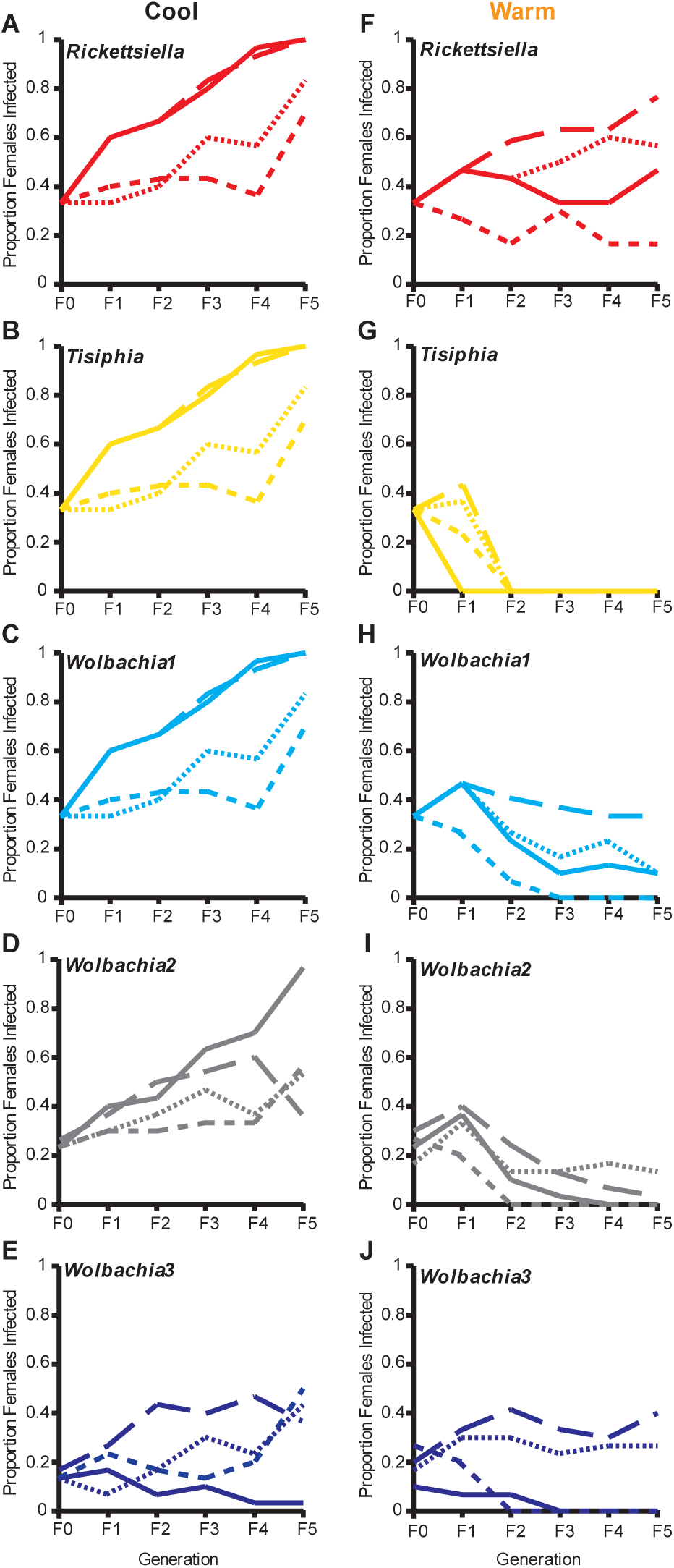
Proportion infection of individual symbionts in laboratory *Mermessus fradeorum* populationsunder A-E) cool (21°C) or F-J) warm (29°C) conditions. Each line represents a replicate population with pro-portion infection depicted for the 30 adult females that were randomly selected to propagatethe next generation.

Given the degree of individuality among the cohort symbionts under warm conditions, we analyzed the infection trajectory of each symbiont separately, finding variable generation, temperature and interactive effects for the five symbionts over the course of the experiment (**Figure 3**). Generation had a significant effect on the infection rate of *Rickettsiella* (χ^2^ = 60.3, d.f. = 5, P < 0.001), *Tisiphia* (χ^2^ = 18.3, d.f. = 5, P = 0.003), and *Wolbachia* 1 (χ^2^ = 12.5, d.f. = 5, P= 0.03) but only a marginal effect on *Wolbachia* 2 (χ^2^ = 10.9, d.f. = 5, P = 0.05) and *Wolbachia* 3 (χ^2^ = 10.4, d.f. = 5, P = 0.06). Temperature also had a variable effect on symbiont spread, significantly influencing the infection rate of *Tisiphia* (χ^2^ = 23.9, d.f. = 1, P < 0.001), *Wolbachia* 1 (χ^2^ = 12.1, d.f. = 1, P < 0.001), and *Wolbachia* 2 (χ^2^ = 25.4, d.f. = 1, P < 0.001), but with a more marginal effect on *Rickettsiella* (χ^2^ = 3.6, d.f. = 1, P= 0.06) and no effect on *Wolbachia* 3 (χ^2^ = 0.7, d.f. = 1, P = 0.41). The interaction between generation × temperature also had a significant effect for every symbiont (R χ^2^ = 31.4, d.f. = 5, P <0.001; T χ^2^ = 139.9, d.f. = 5, P <0.001; W1 χ^2^ = 108.5, d.f. = 5, P <0.001; W2 χ^2^ = 77.2, d.f. = 5, P < 0.001) except *Wolbachia* 3 (χ^2^ = 6.3, d.f. = 5, P = 0.27), indicating that for most symbionts, temperature effects on symbiont infection rate depended on generation.

The divergence between symbiont spread at warm and cool temperatures was driven by the successful spread of most symbionts at cool temperatures but not warm. In cool conditions, *Rickettsiella*, *Wolbachia* 1, and *Tisiphia* frequency increased significantly from the starting infection rate beginning in the F2 generation (P <0.04), *Wolbachia* 2 diverged by the F3 generation (P = 0.004), and *Wolbachia* 3 diverged by the final F5 generation (P = 0.02). In contrast, symbiont population spread was variably diminished in warm conditions. In the most extreme case, *Tisiphia* was completely lost from every warm population within two generations (**Figure 3G**). Both *Wolbachia* 1 and *Wolbachia* 2 also showed significantly reduced infection rates at the warm temperature compared to the cool temperature by the F3 (P <0.001) and F2 (P = 0.005) generations, respectively. Both *Wolbachia* 1 (P = 0.01) and *Wolbachia* 2 (P = 0.01) infection rates in the warm treatment were also significantly lower compared to their starting frequencies at the conclusion of the experiment. *Rickettsiella* frequency only diverged between the two temperature treatments by the final generation (P = 0.002), and *Rickettsiella* ultimately failed to spread significantly from its starting infection rate at warm temperatures due to the extreme variability in its infection rate across the four population replicates (P= 0.31). *Wolbachia* 3 infection also failed to spread in warm populations (P = 0.99), and was not significantly reduced compared to spread at cool temperatures (P = 0.92).

Reduced spread of the symbiont consortium in warm populations was also associated with reduced strength of the feminizing phenotype. Female infection, temperature, and the interaction of maternal infection × temperature all had significant effects on offspring sex ratio (Infection χ^2^ = 426.92, d.f. = 1, P < 0.001; Temperature χ^2^ = 148.55, d.f. = 1, P < 0.001; Inf × Temp χ^2^ = 361.51, d.f. = 1, P < 0.001; **Figure 4A**). Infected females produced significantly female-biased offspring broods compared to uninfected females at both cool (P < 0.001) and warm (P < 0.001) temperatures, but feminization rates were reduced at warm temperatures compared to cool temperatures (P < 0.001). When we restricted the infected category to only spiders that retained *Wolbachia* 1, accounting for the variable loss of symbionts in warm temperatures, results were similar to the general model. Even when *Wolbachia* 1 was retained in warm temperatures, its ability to feminize was reduced relative to cool temperatures (*Wolbachia* 1 infection χ^2^ = 502.76, d.f. = 1, P < 0.001; Temperature χ^2^ = 75.81, d.f. = 1, P < 0.001; W1 Inf × Temp χ^2^ = 288.28, d.f. = 1, P < 0.001; **Figure 4A**).

**Figure 4.**
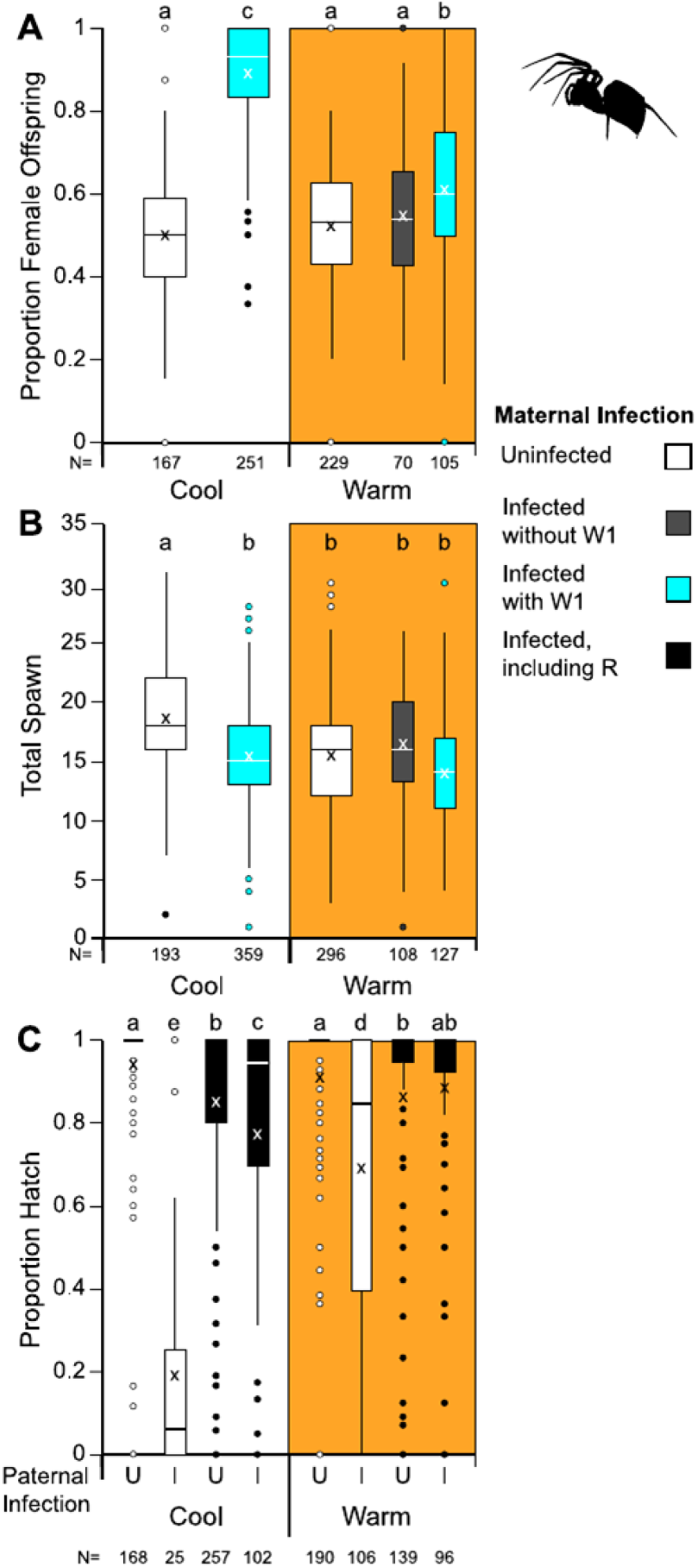
*Mermesst/sfradeorum* offspringA) sex ratio, B) totalspawn (eggs + spiderlings), and C) proportion hatch as a function of maternal infection under cool (21 °C; white background) orwarm (29°C; orange background) conditions in the feminization spread population experiment. For panels A and B, infected spiders are subdivided into those that did (blue) or did not (gray) include the feminizing symbiont, *Wolbachia* 1. For panelC, all infected (I) spiders containedtheCI inducingsymbiont, *Rickettsiella,* and cross type is indicated by paternal infection status on the x-axis. For all panels, sample size isonthex-axis, columns with different lowercase letters are significantly different at α= 0.05

Egg production, another aspect of symbiont-induced phenotypes expected to influence population spread, also differed among populations. Female infection (χ^2^ = 28.8, d.f. = 1, P < 0.001), temperature (χ^2^ = 10.5, d.f. = 1, P = 0.001), and the infection × temperature interaction (χ^2^ = 19.7, d.f. = 1, P < 0.001) all had significant effects on egg production. In cool populations, infected spiders experienced an ∼15% decrease in total egg production as a direct consequence of feminization (**Figure 4B**; P < 0.001), consistent with previous results (Robinson et al. 2026). This fecundity cost disappeared along with feminization in the warm populations, where there was no difference in egg number between infected and uninfected spiders (P = 0.95). As with the sex ratio analysis, follow-up analyses accounting for symbiont loss in warm temperatures yielded similar results to the general model (*Wolbachia* 1 infection χ^2^ = 46.7, d.f. = 1, P < 0.001; Temperature χ^2^ = 16.6, d.f. = 1, P < 0.001; W1 Inf × Temp χ^2^ = 4.8, d.f. = 1, P < 0.03; **Figure 4B**).

Co-infected males induced CI in both cool and warm populations, but uninfected females suffered much higher offspring mortality in cool populations than warm (**Figure 4C**). Cross type (i.e. parental infection status), temperature, and their interaction significantly influenced offspring mortality (Cross Type χ^2^ = 1145.6, d.f. = 3, P < 0.001; Temperature χ^2^ = 22.2, d.f. = 1, P < 0.001; Cross × Temp χ^2^ = 324.9, d.f. = 3, P < 0.001). CI crosses in cool populations were relatively infrequent (25 total from F1 to F5), because the infected females primarily produced daughters, rather than sons that could induce CI. The few males that were produced by infected females, though, induced strong CI, producing significantly more unhatched eggs than compatible crosses (P < 0.001). In warm temperatures where feminization fails, infected males and incompatible crosses were more common (108 total from F1 to F5), and again produced significantly more unhatched eggs than compatible crosses (P < 0.001). However, CI was much weaker in warmer temperatures, with incompatible crosses producing fewer unhatched eggs and higher hatch rates at warm temperatures than cool (P < 0.001).

### Warm temperatures have minimal effects on the spread of the CI symbiont

When we separately evaluated the spread of CI-inducing *Rickettsiella* alone, we found that *Rickettsiella* was consistently able to spread through uninfected spider populations regardless of temperature. Over the course of five host generations, generation significantly influenced infection rate (χ^2^ = 141.9, d.f. = 5, P < 0.001), but neither temperature (χ^2^ = 0.9, d.f. = 1, P = 0.32; **Figure 5**) nor the interaction between generation × temperature (χ^2^ = 7.4, d.f. = 5, P = 0.19) had an effect, and *Rickettsiella* spread equally well regardless of temperature (P= 0.59). *Rickettsiella* infection rate began diverging significantly from the starting infection rate by the F3 generation under both cool (P < 0.001) and warm (P = 0.003) conditions.

**Figure 5.**
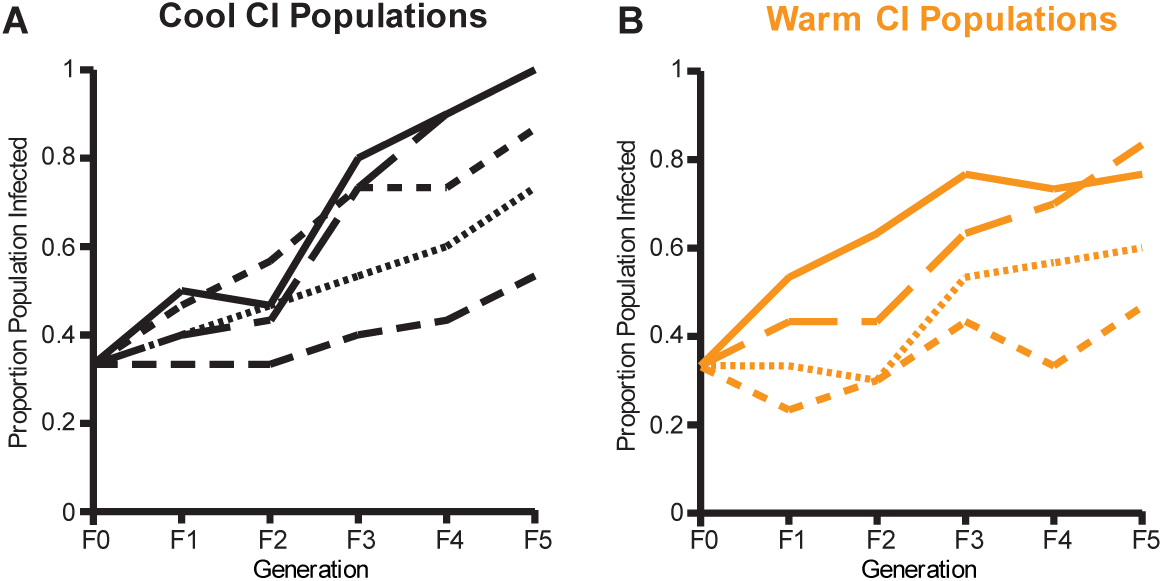
Proportion infection of laboratory *Mermessus fradeorm* populations under A) cool (21°C) or B) warm (29°C) conditions in the CI spread populations. Each line represents a replicate population, with proportion infection depicted for the 30 adult females that were randomly selected to propagate the next generation. Infected spiders were infected with only *Rickettsiella*.

These effects were largely driven by incompatible CI crosses between *Rickettsiella* males and uninfected females (Cross χ^2^ = 589.78, d.f. = 3, P < 0.001; Temp χ^2^ = 53.43, d.f. = 1, P < 0.001; Cross × Temp χ^2^ = 51.67, d.f. = 3, P < 0.001), which produced significantly higher amounts of unhatched eggs compared to compatible crosses at cool (P < 0.001) and warm temperatures (P < 0.001; **Figure 6C**). As with the CI mortality in the feminized populations, *Rickettsiella* CI was significantly weaker at warm temperatures compared to cool temperatures (P < 0.001), which resulted in comparatively higher rates of successfully hatched spiderlings from lethal CI crosses at warm temperatures. In contrast to the feminized populations, offspring sex ratios in the CI populations were not significantly affected by maternal infection (χ^2^ = 1.95, df = 1, P = 0.16), temperature (χ^2^ = 1.59, d.f. = 1, P = 0.21), or their interaction (χ^2^ = 0.006, d.f. = 1, P = 0.94; **Figure 6A**). However, total offspring production was significantly affected by parental infection (χ^2^ = 8.2, d.f. = 1, P = 0.004) and temperature (χ^2^ = 21.3, d.f. = 1, P < 0.001), but not their interaction (χ^2^ = 2.2, d.f. = 1, P = 0.14; **Figure 6B**). While at cool temperatures egg production did not differ between infected and uninfected females (P = 0.56), under warm conditions *Rickettsiella* infected females produced more eggs than uninfected ones (P = 0.02). Egg production was also generally reduced at warm temperatures compared to cool conditions for both uninfected (P <0.001) and *Rickettsiella*-infected females (P < 0.001).

**Figure 6.**
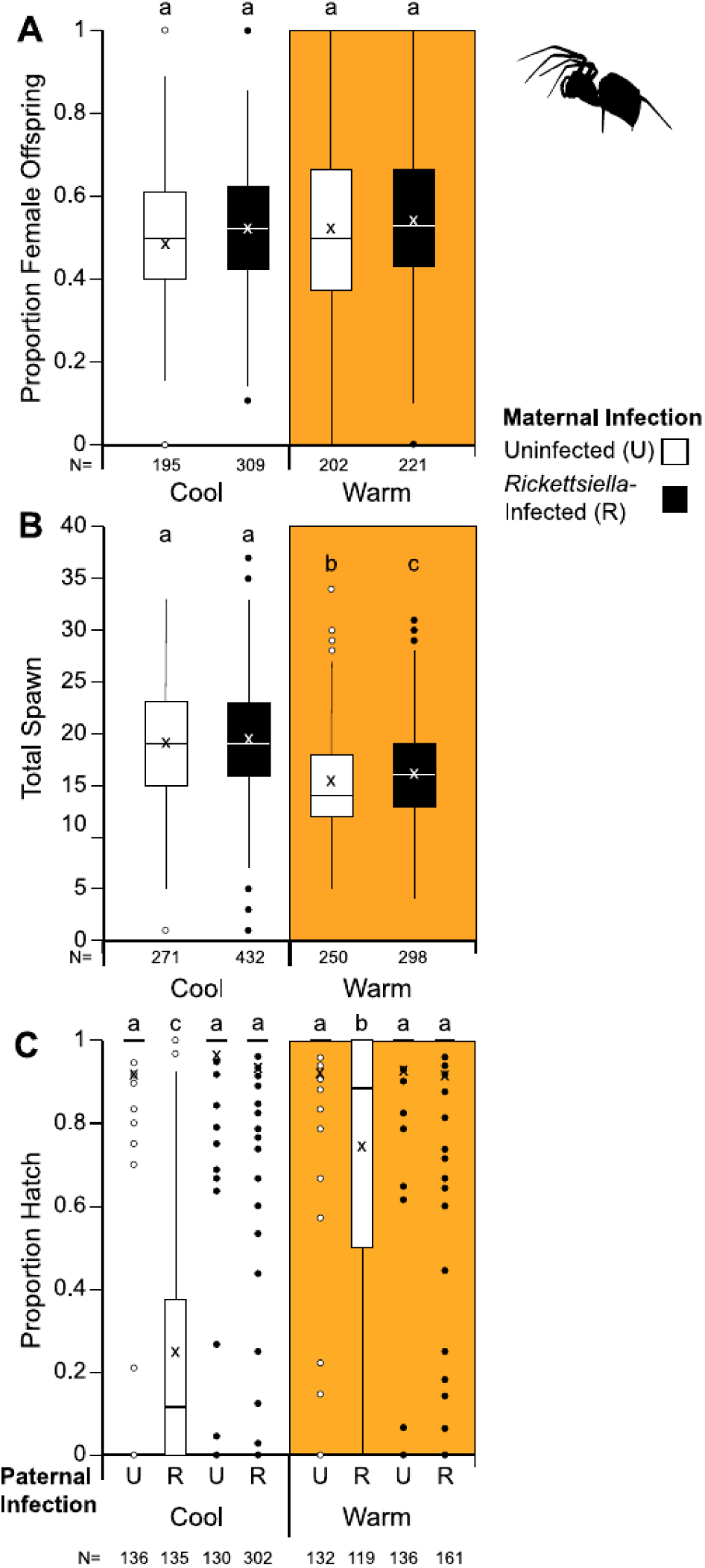
*Mermessus fradeorum* offspring A) sex ratio, B) total spawn (eggs + spiderlings), and C) proportion hatch as a function of maternal infection with Rickettsiella under cool (21°C; white background) or warm (29°C; orange background) conditions in the Cl spread population experiment. For proportion hatch data (panel C), cross type is indicated by paternal infection status on the x-axis. For all panels, sample size is on the x-axis, columns with different lowercase letters are significantly different at α = 0.05

## Discussion

Our study provides empirical demonstration that two separate reproductive manipulations, CI and feminization, can propel the spread of a heritable bacterial community through a host population. Despite clear theoretical predictions that manipulative symbionts will increase in frequency (Caspari and Watson 1959; J. Engelstädter and Telschow 2009; Hatcher 2000; O’Neill et al. 1997; Perlman et al. 2008; Turelli 1994), controlled experimental studies that document the phenomenon are surprisingly scarce (Harris et al. 2010; Schmidt et al. 2017; Xi et al. 2005) particularly for feminization (Miyata et al. 2024). Under cool conducive conditions, where symbiont vertical transmission rates were high and symbiont-induced manipulative phenotypes were strong, both the CI-inducing *Rickettsiella* and the feminizing *Wolbachia* were able to increase to more than 80% of the host population within 5 generations. Further, co-infecting symbionts that did not exert discernable phenotypic effects on their host, such as *Tisiphia*, were likewise able to spread through the host population when under permissive environmental conditions.

Under warmer conditions, however, chaos ruled. All members of the symbiont community experienced reduced vertical transmission rates, ranging from a subtle decrease, in the case of *Rickettsiella*, to complete extinction in the case of *Tisiphia* (see also Mackevicius-Dubickaja et al. 2026). As a consequence, the symbiont community in *M. fradeorum* shifted from a stable consortium to a mix-and-match assortment of bacteria. Both CI and feminization phenotypes were weakened as well: For *Rickettsiella*, the combination of strong transmission and weak CI was sufficient to permit spread when *Rickettsiella* was the only symbiont present. As a member of the dissolving feminizing consortium, however, *Rickettsiella* was only able to maintain a steady infection rate. In contrast, the feminizing *Wolbachia* did not have a sufficient combination of transmission and manipulation to maintain itself (Brenninger et al. 2025), and declined in the host population. Likewise, co-infecting non-manipulative symbionts were unable to effectively hitchhike and uniformly failed to spread.

Despite prior evidence of tradeoffs in titer among co-infecting symbionts in *M. fradeorum* (Mackevicius-Dubickaja et al. 2025), the present study emphasizes that when environmental conditions are permissive, the co-infection appears to be stable and mutually beneficial for the participating microbes. The feminization caused by *Wolbachia* 1 drives spread through the host population, despite the unavoidable fitness costs imposed by this manipulation (Robinson et al. 2026). *Rickettsiella* contributes to consortium spread as well, albeit more subtly, by weaponizing the occasional males produced by imperfect feminization to kill uninfected offspring via CI. As feminization spread and males became rare in the population, matings between these infected, sabotaged males and uninfected females became more common, which accelerated spread of the symbiont consortium near the end of the experiment.

The contributions of the other symbionts to the consortium under cool conditions remain equivocal. As has been shown previously (Mackevicius-Dubickaja et al. 2025), the presence of *Wolbachia* 3 strengthened *Wolbachia* 1 feminization, resulting in 93.6±0.9% female offspring for spiders infected with RTW123, compared to 86.8±1.6% for spiders infected with RTW12, and 86.3±2.1% for those with RTW1 (**Supplemental Figure S2**). However, this boost in feminization did not result in an evident advantage for the RTW123 consortium over the RTW12 or RTW1 symbiotypes and the overall proportion of these three symbiotypes relative to one another under cool conditions remained remarkably consistent throughout the experiment (**Supplemental Figure S1**). It is possible that the benefits of stronger feminization in RTW123 females versus the production of more CI-inducing males in RTW12 and RTW1 might have balanced each other out. *Tisiphia* was present in all feminized spiders under cool conditions, which precludes any direct insights about its role in the consortium. It may simply be the most fortunate hitchhiker, or it might be contributing to the community in ways that have yet to be determined. Previous work (Mackevicius-Dubickaja et al. 2026) has suggested that loss of *Tisiphia* may have a destabilizing effect on the community, which would be consistent with our observations under warm conditions in the present study. It is possible that the non-manipulative members of the consortium contribute to community stability in other less obvious ways, perhaps by improving symbiont transmission rates (Rock et al. 2018) or mitigating host fitness costs (Doremus and Oliver 2017; Xie et al. 2016), rather than acting as passive hitchhikers.

Under warm conditions, however, vertical transmission failure was rampant. Along with the complete loss of *Tisiphia* by the F2 generation, all three *Wolbachia* strains experienced dozens of transmission failures over the course of the experiment, resulting in extinction in at least some populations (**Figure 3**). Interestingly, *Wolbachia* 3, which is the only symbiont of the consortium to suffer incomplete transmission at both warm and cool temperatures (Mackevicius-Dubickaja et al. 2026), and which started at the lowest proportion of the populations, seemingly weathered the warm temperatures the best of the *Wolbachia* strains by maintaining a stable infection rate in two of the four warm temperature populations. While *Wolbachia* 3 may be the more thermally robust of these *Wolbachia*, it’s lack of host effect and generally incomplete transmission likely inhibits its overall spread. Only *Rickettsiella* retained consistent transmission in warm conditions, suffering only 11 instances of apparent loss over the course of the experiment, mostly from matrilines that had already lost their other symbionts. The apparent robustness of *Rickettsiella* extends to other environmental threats, including antibiotics, and this symbiont is generally the last symbiont to be lost under stressful conditions (Mackevicius-Dubickaja et al. 2025). Future genomic analyses may provide some insight into why this *Rickettsiella* strain fares better than its colleagues under stressful conditions. Similarly, gene expression analyses will provide more context to the abrupt transmission failure of symbionts like *Tisiphia* and the variability in temperature sensitivity across the more closely related *Wolbachia* strains.

The symbiont spread observed under warm experimental conditions generally aligns with patterns observed for the *M. fradeorum* symbiont community in fields in the East central United States. In these fields, virtually all *M. fradeorum* spiders (>99%) are infected with *Rickettsiella* (Rosenwald 2020; Rosenwald et al. 2020; M. Doremus and J. White, unpublished data), while a stable minority (∼20%) of the population host the complete five-member consortium, with a further subset of spiders hosting a mixture of other subsidiary symbiotypes (J. White, unpublished data). Our results suggest that environmental factors, most notably temperature, reinforce this stable multiple-member symbiont community, with seasonal swings between cool conducive conditions potentially encouraging the spread of the feminizing consortium, while warm disruptive conditions winnow away all but *Rickettsiella*. Of course, our uniform laboratory conditions reflect a more severe thermal treatment than that experienced by spider and symbiont in nature, as both temperature fluctuations and spider thermoregulatory behaviors likely mitigate thermal stress in field settings. This caveat may explain why *Tisiphia* was lost from experimental populations despite being the second most common symbiont in field *M. fradeorum* populations (Rosenwald et al. 2020; Rosenwald 2020, M. Doremus and J. White, unpublished data). Symbiont frequencies in natural settings are further complicated by the multi-generational effects of temperature on symbiont stability and host phenotype (Mackevicius-Dubickaja et al. 2026), and the multivoltine life cycle of the spider host, which has overlapping generations potentially developing under different thermal conditions.

Our understanding of how environmentally sensitive heritable symbionts spread in a world of changing temperatures remains limited, with scant studies empirically testing how environment shapes symbiont population spread. Using experimental populations held at different temperature conditions, we found that both external and internal environmental contexts shape the efficacy of symbiont spread, with elements like temperature and co-infection both contributing to the spread or loss of symbionts. We also found substantial variation in symbiont thermal susceptibility, with some symbionts demonstrating efficient spread despite weakened host effects, while others were doomed by reduced vertical transmission rates. Our results suggest that temperature is a primary force promoting the stable coexistence of multi-member symbiont communities and may help prevent symbiont-induced sex ratio distortion from collapsing host populations.

## Acknowledgements

We thank Kaitlin Butler, Andy Fajardo, Veronica Kegley, Allie Reagan, Emmett Roberson, Becca Robertson, Laura Rosenwald, and Ellen Williams for invaluable assistance in the lab, and the many spiders that lived and died in the lab as part of this study. We thank Becca Robertson for the spider silhouette used in the Figures.

