## Supplemental material for "Warm temperature impedes the spread of a heritable manipulative symbiont community in spider populations"

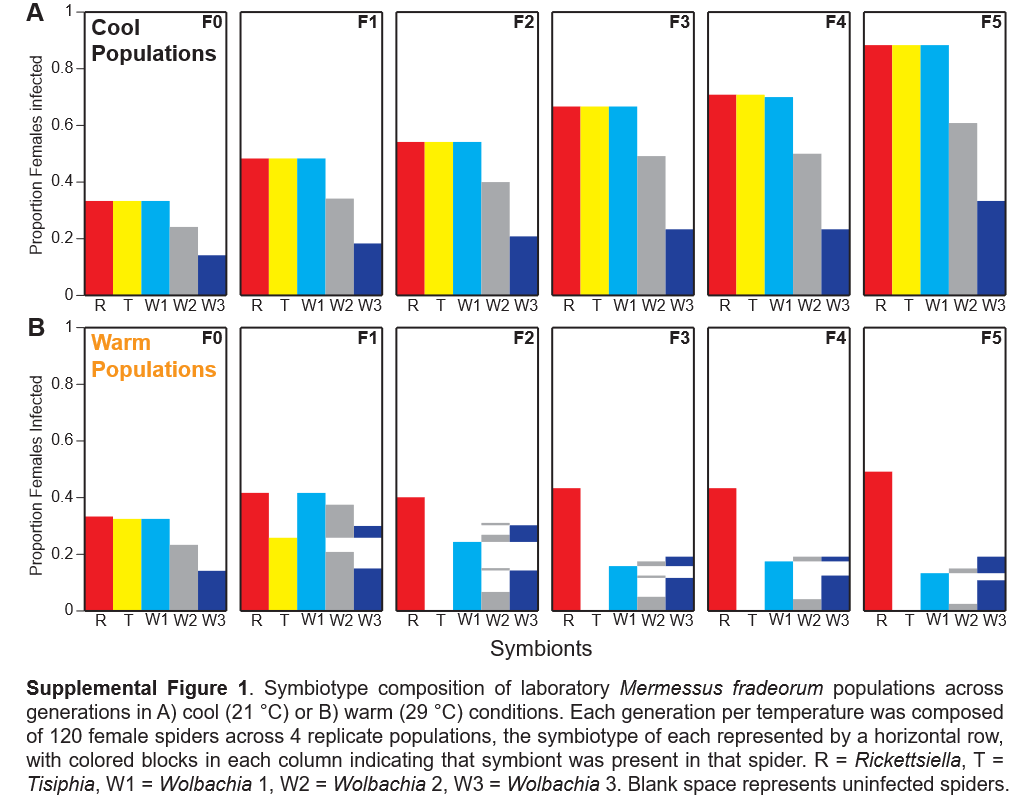

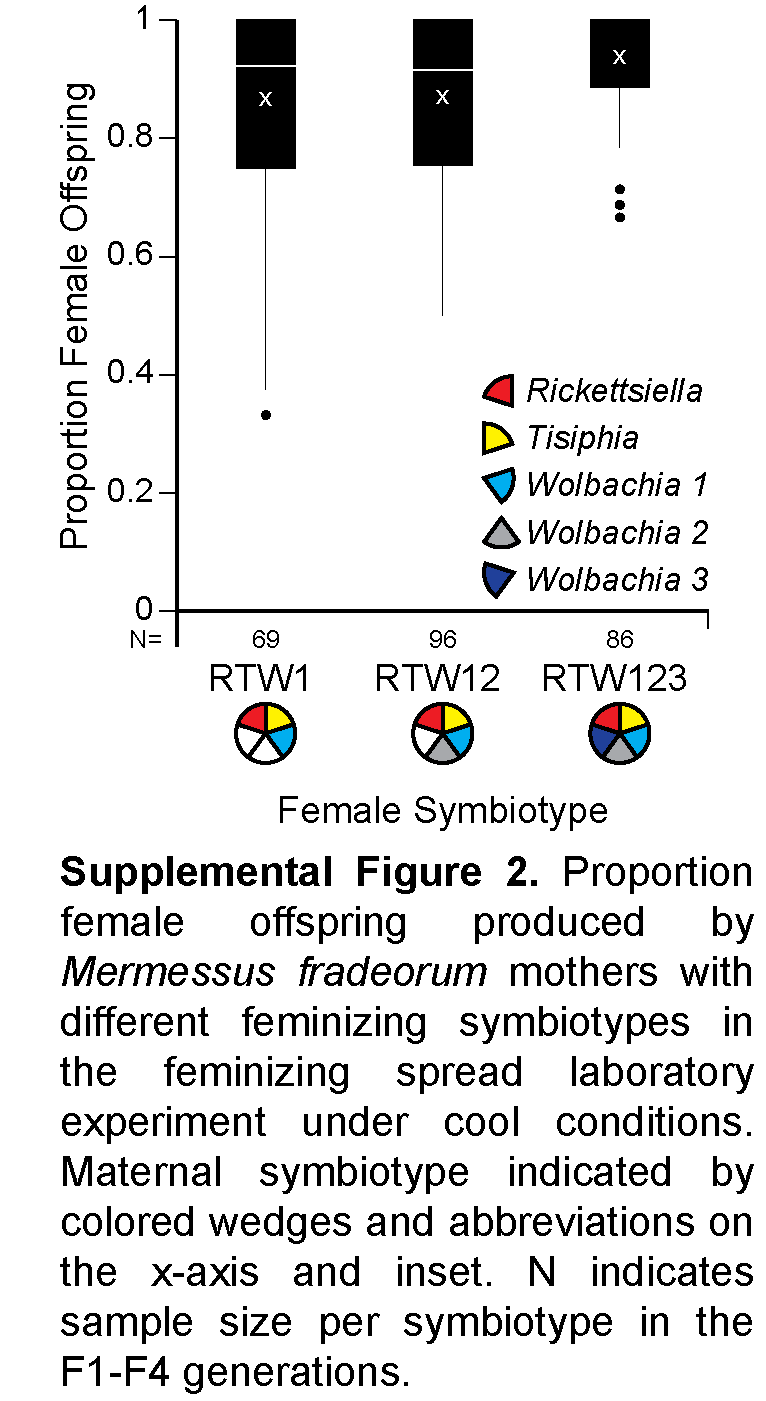

**SI Table 1** Primer pairs used in this study, with references provided for previously published primers for each symbiont used in diagnostic PCR.

| **Target** | **Target gene** | **Primer name** | **Sequence** | **Annealing Temp (°C)** | **Amplicon length (bp)** | **Citation** |
| --- | --- | --- | --- | --- | --- | --- |
| *Rickettsiella* | *16S rRNA* | RLA16s F1 | CAGTAAARRTTTCGGYCTTTAYGGG | 56 | 532 | Duron et al. 2016 |
|  |  | RLA16s R1 | CAAACCTAGTCAACCACCTACACG |  |  |  |
|  | *recA* | recA_345F | ACAACCTGATACTGGCGAGC | 60 | 166 | Proctor et al. 2024 |
|  |  | recA_510R | CGACATTAATCGCGCTTGCA |  |  |  |
| *Tisiphia* | *16S rRNA* | RicklongF | ACGTGGGAATCTACCCATCA | 60 | 530 | Curry et al. 2015 |
|  |  | RicklongR | TAGCCTAGATGACCGCCTTC |  |  |  |
| *Wolbachia* 1 | *wsp* | wsp1_36F | ACAAAAGCATCAGGTCAAGAAAAT | 60 | 167 | Mackevicius-Dubickaja et al. 2025 |
|  |  | wsp1_201R | CATCTGCAGCATTGGTATCATTT |  |  |  |
| *Wolbachia* 2 | *wsp* | wsp2_93F | GCAAGGCAACAAATAAAGACAAGG | 60 | 163 | Mackevicius-Dubickaja et al. 2025 |
|  |  | wsp2_255R | ACATTTGTCTCAGCAGCAGC |  |  |  |
| *Wolbachia* 3 | *wsp* | wsp3_270F | ACCGCTGTGAATGATCAAAACA | 60 | 162 | Mackevicius-Dubickaja et al. 2025 |
|  |  | wsp3_431R | ATCGTTATTAGTTGATGTTGTTGCTT |  |  |  |
| Spider | CO1 | lco1490 | GGTCAACAAATCATAAAGATATTGG | 53 | 650 | Folmer et al. 1994 |
|  |  | hco2198 | TAAACTTCAGGGTGACCAAAAAATCA |  |  |  |

Curry, M. M., Paliulis, L. V., Welch, K. D., Harwood, J. D., & White, J. A. (2015). Multiple endosymbiont infections and reproductive manipulations in a linyphiid spider population. *Heredity (Edinb), 115*(2), 146-152. doi:10.1038/hdy.2015.2

Duron, O., Cremaschi, J., McCoy, K.D. (2016). The high diversity and global distribution of the intracellular bacterium *Rickettsiella* in the polar seabird tick *Ixodes uriae*. *Host Microbe Interact* 71(3): 761-770. doi: 10.1007/s00248-015-0702-8

Folmer, O., Black, M., Hoeh, W., Lutz, R., & Vrigenhoek, R. (1994). DNA primers for amplification of mitochondrial cytochrome c oxidase subunit 1 from diverse metazoan invertebrates. *Mol Marine Biol Biotech, 3*, 294-299.

Mackevicius-Dubickaja, V., Gottlieb, Y., White, J. A., & Doremus, M. R. (2025). *Wolbachia* feminises a spider host with assistance from co-infecting symbionts. *Environ Microbiol, 27*(7), e70149. doi:10.1111/1462-2920.70149

Proctor, J. D., Mackevicius-Dubickaja, V., Gottlieb, Y., & White, J. A. (2024). Warm temperature inhibits cytoplasmic incompatibility induced by endosymbiotic *Rickettsiella* in spider hosts. *Environ Microbiol, 26*(9), e16697. doi:10.1111/1462-2920.16697
